# Shared generative rules of locomotor behavior in arthropods and vertebrates

**DOI:** 10.1101/031716

**Authors:** Alex Gomez-Marin, Efrat Oron, Anna Gakamsky, Dan Valente, Yoav Benjamini, Ilan Golani

## Abstract

The discovery of shared behavioral processes across phyla is an essential step in the establishment of a comparative study of behavior. We use immobility as an origin and reference for the measurement of locomotor behavior; speed, direction of walking and direction of facing as the three degrees of freedom shaping fly locomotor behavior; and cocaine as the parameter inducing a progressive transition in and out of immobility. In this way we expose and quantify the generative rules that shape part of fruit fly locomotor behavior, bringing about a gradual buildup of freedom during the transition from immobility to normal behavior and a precisely opposite narrowing down during the transition into immobility. During buildup the fly exhibits enhancement and then reduction to normal values of movement along each degree of freedom: first, body rotation in the horizontal plane, then path curvature and then speed of translation. Transition into immobility unfolds by narrowing down of the repertoire in the opposite sequential order, showing reciprocal relations during both buildup and narrowing down. The same generative rules apply to vertebrate locomotor behavior in a variety of contexts involving transition out and into immobility. Recent claims for deep homology between the arthropod central complex and the vertebrate basal ganglia provide an opportunity to examine whether the generative rules we discovered also share common descent. Neurochemical processes mediating the buildup of locomotor behavior in vertebrates could guide the search for equivalent processes in arthropods. The measurement methodology we use prompts the discovery of candidate behavioral homologies.

**Significance Statement:** Do flies and mice share the same behavior? By defining immobility as an intrinsic reference point for locomotor behavior we show that the rules that generate the transition from immobility to full blown normal behavior, and from full blown behavior to immobility are shared by fruit flies and mice. These rules constitute a much desired aim of evolutionary biology: the discovery of behavioral homologies across distant phyla. The methodology we use facilitates the discovery of cross-phyletic behavioral homologies, shedding light on the problem of the evolution of behavior.

## Introduction

The establishment of homologies is an indispensable goal in evolutionary biology. In pre-Darwinian comparative anatomy, a homologue has been defined as "The same organ in different animals under every variety of form and function"^1^. Based on this definition, anatomists have compared skeletons using validated distinctions such as a forelimb, a humerus, and a radius, and compared brains using validated structures such as the thalamus, cortex, and striatum. These structures acquired their identity and validity as homologues by demonstrating that they occupied the same relative position, and had the same connectivity across a wide array of taxonomic groups^2^ and structures sharing the same name and the same morphogenetic history. The validity of these structures has been indispensable for establishing a rigorous science of anatomy. Similarly, the comparative study of behavior requires the identification of distinct elementary processes, much like skeletal segments and neural structures, which could be established across a wide variety of taxonomic groups on the basis of their connectivity^3^, and their moment-to-moment generative history.

The accumulation of detailed descriptions of arthropod and vertebrate movement makes the issue of shared principles of organization in the behavior of these taxonomic groups increasingly accessible for comparison. An opportunity for such comparison is offered by the report that, when treated with the dopamine reuptake inhibitor cocaine, *Drosophila melanogaster* performs a sequence of stereotyped behavior patterns, including locomotion and circling, that lead in and out of immobility, apparently similar to the sequence observed in rodents^4,5^. This led the researchers to suggest that the behavior was homologous in the two phyla. That same behavior has, however, been portrayed as aberrant, unusual, and uncontrolled^6^ by others, whose description highlighted impairment in the functionality of the behavior.

Here we analyze the structure of this behavior in its own right, while also using a strategy and tools enabling a cross-phyletic comparison: by focusing on structure, we derive from the fly’s seemingly aberrant behavior the generative rules^7,8^ that shape a substantial component of both arthropod and vertebrate locomotor behavior. Studying the morphogenesis of cocaine-induced fruit fly behavior, we implement a strategy and tools that seek to reveal those generative rules with the potential to become universal with the very first description of a newly-studied behavior.

We suggest that cross-phyletic behavioral homologies have hardly been documented so far because the seed for obtaining a phylogenetic perspective (or for missing it) is sown in the initial measurement phase by the choices made by the observer. These choices determine from the start the potential for the universality of the description. Since they are inescapable, they might as well be made deliberately, on the basis of one’s particular aims. Our aim, of establishing a rigorous comparative study of behavior, requires the discovery of the generative rules that will have the potential of defining a *bona fide* behavioral homology. The choices we made are derived from this aim (other aims may justify other choices). While any selection of variables might be informative, only the key variables that are actively managed by the fly have the potential of defining cross-phyletic generative rules. Conversely, kinematic quantities that prove universal across phyla are more likely to represent perceptual quantities that are actually managed by the brain. A judicious selection of the key variables that describe the behavior is therefore necessary for uncovering cross-phyletic homologies.

One strategy in deciphering the organization of an anatomical structure is to follow the process of its morphogenetic differentiation from inception to the full blown form. The transition from simple to complex provides a view that cannot be obtained by studying the final product alone. We therefore focus on studying fly behavior in a situation involving the differentiation from simple to complex, and the subsequent decay from complex to simple, teasing apart in this way the particulate processes that add on top of each other to compose full-blown behavior^9,10^ (or are eliminated in sequence until reaching full decay).

Recent claims for a deep homology between the arthropod central complex and the vertebrate basal ganglia^21^ provide an opportunity to examine whether the generative rules we discovered can be supplemented with a historical perspective. If cocaine induced behavior, which is mediated by these centers, is the same in the two phyla, then the claim for a behavioral homology would be supported by both a claim for common generative rules and a claim for common descent. Moreover, the shared generative rules can provide a specification of the demand^9,11^ on the neural activity and network connectivity within and between substructures of the central complex and the basal ganglia. For example, the buildup of a vertebrate’s locomotor repertoire, which has been recently attributed to dopaminergic feedforward loops operating in the basal ganglia^12,13^, can guide a study of the relations between the arthropod transition out of immobility and the central complex.

## Results

### Narrowing down of the path’s spatial spread and its build up to spatially spread normal behavior

Figure 1 presents the path traced by a single fly walking in the arena throughout a 90 min session. Upon cocaine administration, a complex dynamics of movement leads to immobility (marked by the red dot), followed by full recovery of movement (Fig. 1). The path leading to immobility is colored in blue, and the path leading out of it is colored in green. The path traced in space (Fig. 1A) unfolds in time (Fig. 1B) highlighting immobility as the origin to which motion converges and from which it unfolds. The fly first traces relatively straight paths, which become increasingly more curved culminating in immobility (Fig. 1C). Transition out of immobility starts with highly curved paths involving many tiny circles, followed by increasingly straighter paths (Fig. 1D). The progressive narrowing down of the path into immobility (Fig. 1E) and its progressive buildup back to normal (Fig. 1F) is quantified for all flies.

**Figure 1.**
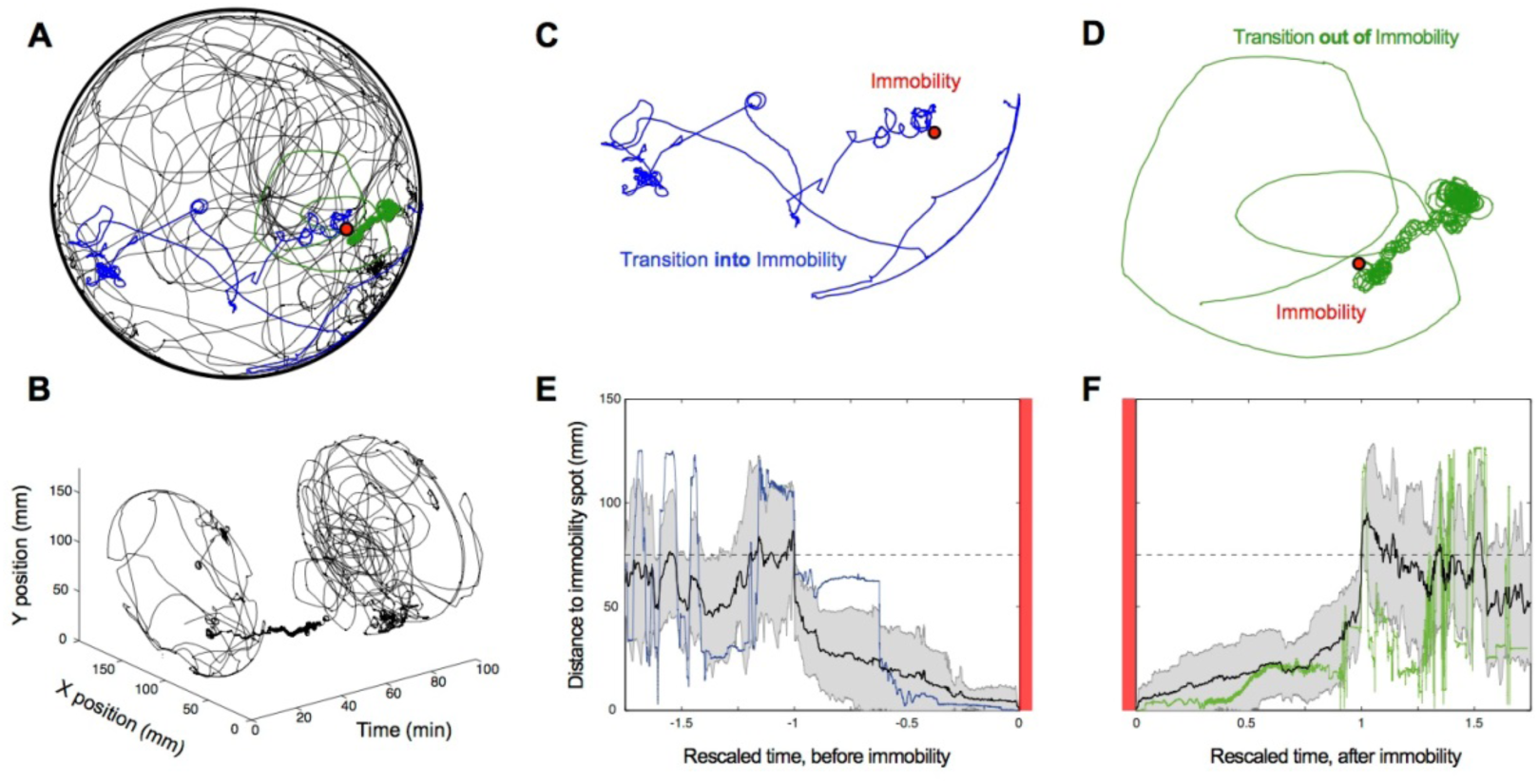
The locomotor path of cocaine treated flies narrows down into immobility and builds up to spread-out normal walking behavior. (**A**) **A** fly’s path for the entire **90** min session in a circular arena. Red dot indicates location of immobility. Blue path depicts transition into immobility, and green transition out of it. **(B)** Same path as in **(A)** unfolded in time. (C) The transition into immobility is marked by the performance of straight paths, and then by increasingly more curved paths, narrowing down the spatial spread of the animal’s path. (**D**) The transition out of immobility is marked by the performance of curved, then increasingly straighter paths, building up the spatial spread of the animal’s path. (**E**) By rescaling time in reference to immobility (marked by the vertical red lines), we demonstrate the narrowing of the path for all flies during transition into immobility, and (**F**) the buildup of the path for all flies during transition out of immobility.

### Narrowing down of the fly’s locomotor repertoire and its build up to a normal repertoire

The three degrees of freedom of locomotor behavior studied are speed, direction of walking, and body orientation (Fig. S1 and Methods). Using extended immobility as a reference, we trace the behavior that preceded it (Fig. 2), starting with the inflow of cocaine into the arena, and ending with full recovery of the fly, as marked by the disappearance of high rotation in place (extensive changes in body orientation at low translational speed) and the performance of straight paths (high speed at low curvature). The entire dynamics in terms of speed (Fig. 2A), curvature (Fig. 2B), and rotation (Fig. 2C) are illustrated for a single fly. Immobile for 10 minutes (gray shaded area), recovery from immobility started with very fast whole-body rotations in place. Each diagonal line in magenta stands for a full 360 degree rotation. The fly performed approximately 50 full rotations in 10 minutes with almost zero translational velocity and extremely high path curvature. The high frequency of rotations gradually decreased, as did the path curvature (Fig. S2). Then the animal resumed normal forward progression involving relatively straight paths and high velocity. Transition into immobility started with normal locomotion marked by high speed, low curvature, and absence of extensive body rotations. Next, we observed bursts of high velocity followed by medium and then high curvature, which concurred with the setting in of rotations at very low translation speeds, culminating in immobility.

**Figure 2.**
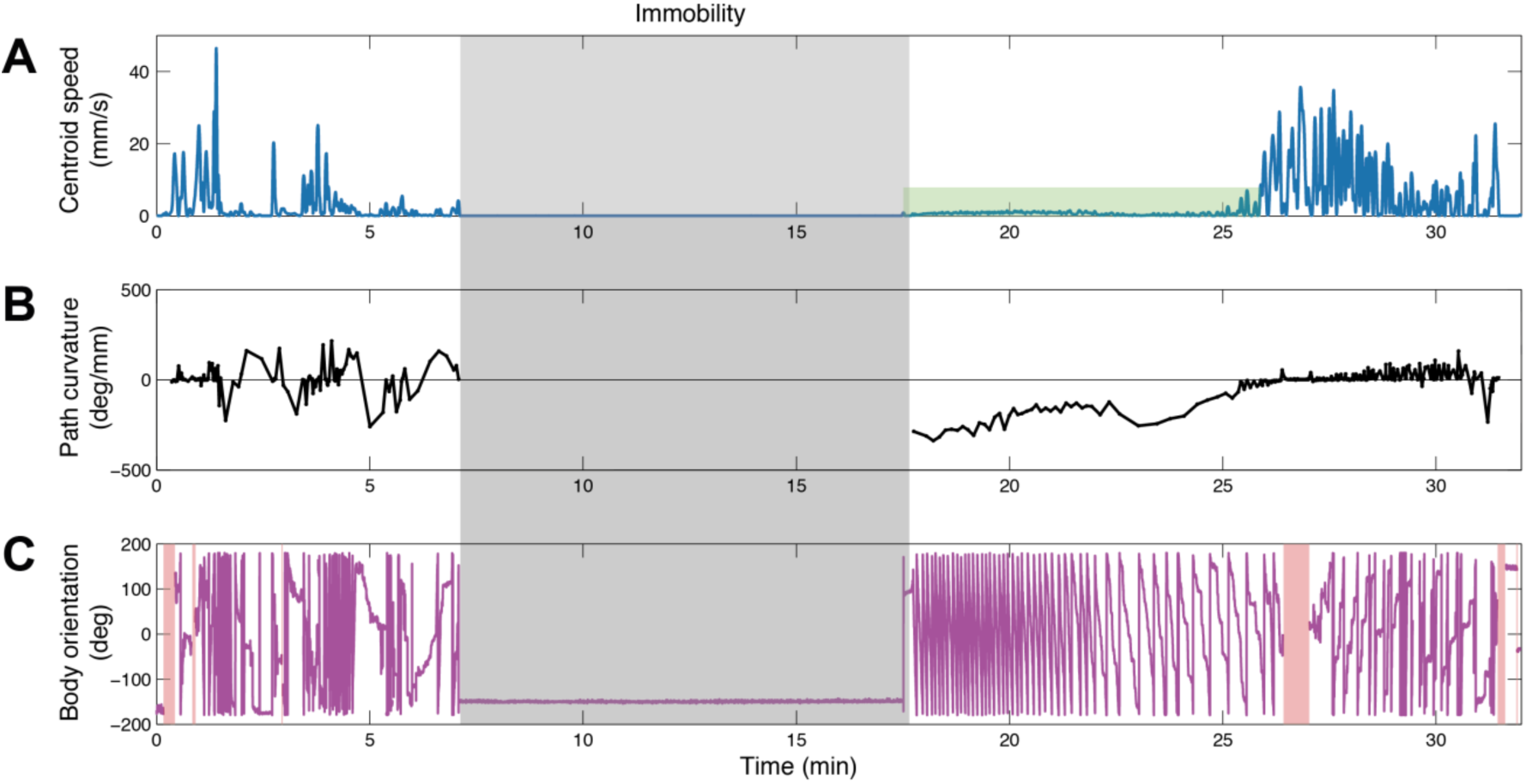
Representative moment-to-moment dynamics of the three kinematic degrees of freedom of a single fly for an entire session. The shaded area marks the period of immobility, which is used as a reference for the events that precede and follow it. (**A**) The session starts with bursts of speed that progressively decrease towards zero. Following a 10 minute period of complete immobility, very low speed is then followed by normal speed. The green shaded area highlights small but non-zero velocity components at high-curvature during rotation in place (shown in detail in Fig. S2). (**B**) Straight path is followed by bursts of high curvature until immobility, from which the fly resumes its movement with very high curvature (of the order of a 360 degree turn in a millimeter) monotonically decreasing to straight paths again. (**C**)Extensive body rotations precede and follow immobility, proceeding from low to high frequency, and from high to low frequency. Red shaded areas represent time segments when the fly touches the walls of the arena and body orientation is not tracked. Each diagonal line represents a 360 degree body rotation. Overall, the session starts with extensive translation, then increasing curvature accompanied by frequent body rotations, leading into immobility. Following immobility, extensive rotation in place concurring with very high path curvature, is followed by forward progression along straight paths.

We can summarize the moment-to-moment dynamics observed as the following sequence: predominance of translation, then high curvature, and finally extensive rotation in place, ending in immobility, from which the same sequence is performed in reverse. Forward translation is thus eliminated from the fly’s repertoire first, and rotation last, in the transition into immobility (movie S1); while rotation is added to the repertoire first, and forward translation last, in the transition out of immobility (movie S2). Animations in time of the progressive narrowing down of degrees of freedom for movement (movie S3) and of progressive buildup (movie S4) clearly represent the dynamics of the phenomenon under study.

### The direction in which a fly walks and the direction it faces alternate in who leads and who follows

Flies can walk in different directions while keeping their body orientation fixed, or else facing in any other direction while proceeding in a specific direction. In intact flies, during walking, the direction of progression changes first, and the direction the fly faces then converges to the new direction set by progression, with facing lagging behind by a small angular interval that is quickly closed. The same order of leading and following is exhibited under cocaine administration, but facing direction lags behind by a much larger angular interval and closing the interval takes a much longer time^14^. During the stage of transition out of immobility (Fig. 3A), as flies rotate along highly curved paths, the directions of progression and of facing tend to converge to the same values (Fig. 3B-C). The further away from immobility, the coupling between these two degrees of freedom becomes progressively less tight (Fig. 3C-D). Furthermore, first the direction of progression leads while body orientation follows; and conversely, later, as the fly regains its freedom of movement away from immobility, body orientation leads and direction of progression follows (Fig. 3E). In other words, the angular interval between the direction of progression and body orientation is modulated dynamically with respect to immobility and actively managed in two opposite ways.

**Figure 3.**
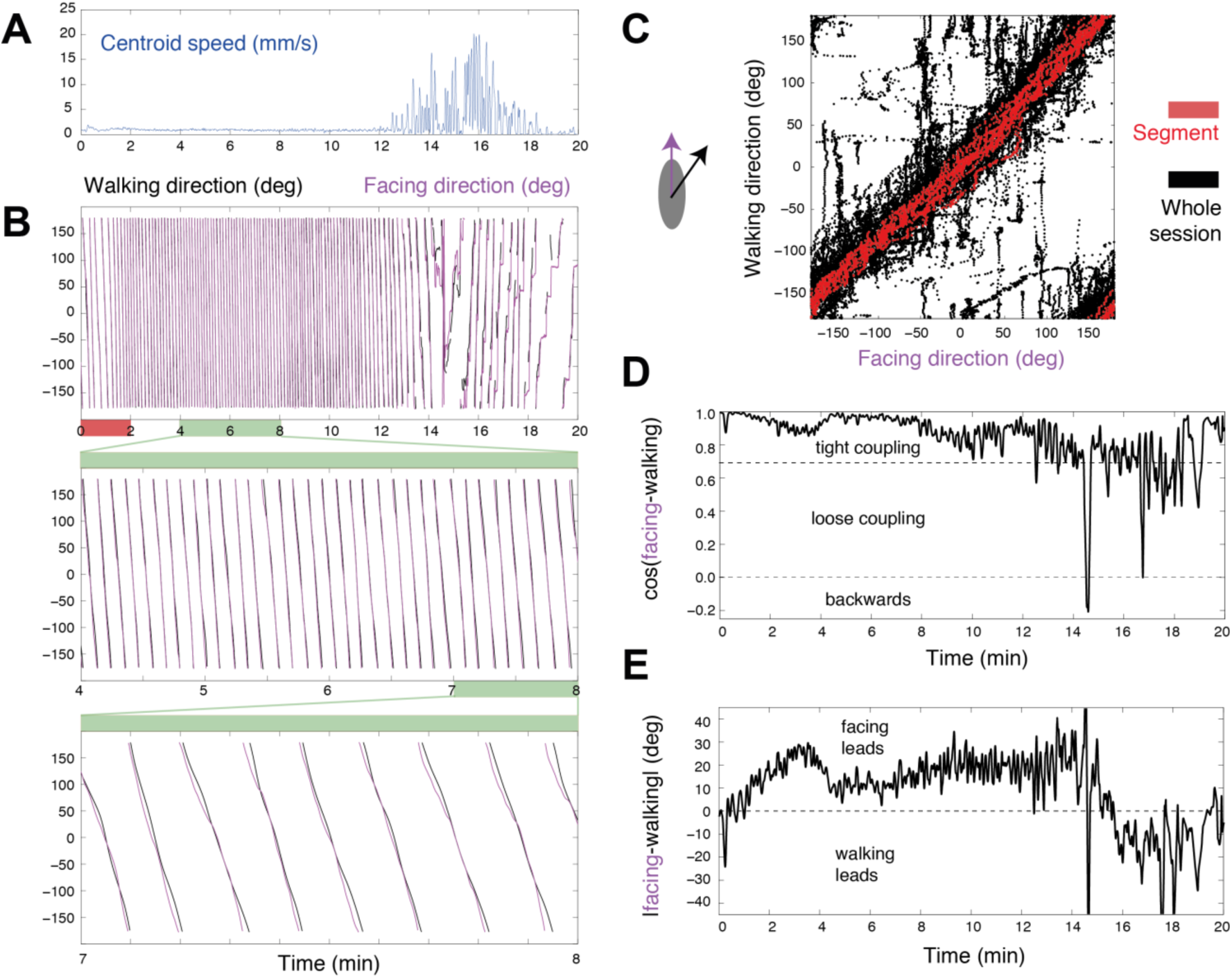
Active management of walking and facing directions as reflected in the dynamics of curvature and rotation. (**A**) Time course of centroid speed during transition out of immobility. (**B**) Walking and facing directions during intense rotation in place and high-curvature dynamics. A two-minute segment of the top plot is amplified in the middle plot; and a one-minute segment of the middle plot is amplified in the bottom plot, where the tight, but not perfect, coordination between both angles (facing and walking directions) can be appreciated. (**C**) Phase-plot of facing-walking direction value combinations across an entire session (black), and selected segment showing that close to immobility the coordination is tight (red). (**D**) Cosine of the difference of facing and walking angles, progressively decreasing from 1 (zero difference), to 0.7 (45 degree difference), and reaching below 0 (more than 90 degree difference), quantifying the transition from tighter to looser coupling of rotational and curvature degrees of freedom as the fly transitions out of immobility. (**E**) Quantification of the difference in angle between the degree of freedom that leads and the one that follows. Facing leads when close to immobility, with walking direction lagging behind a few degrees. Walking direction leads at later stages.

### Generative rules of rotational-translational locomotor behavior in and out of immobility

Having discussed the relationship between curvature and rotation, we concentrate on the relationship between translation (T) and rotation (R). Since the timescale we examine comprises the entire session dynamics (which can last for more than an hour), we next calculate the cumulative translation by integrating velocity in time and cumulative rotation, by unwrapping the body angle (Fig. 4A). After smoothing the cumulative measures (see Methods) we calculate their derivative, and thereby obtain the changes in rotation and translation for the entire session (Fig. 4B). In order to quantify the sequence in which they unfold, we again use immobility as a reference and measure the global peaks of activity before and after immobility, along each of the two degrees of freedom. This procedure reveals that before immobility the maximal peak of translation (T*) is exhibited before the maximal peak of rotation (R*). After immobility, it is the maximal peak of rotation that is exhibited before the maximal peak of translation. To quantify the relative order of global peaks, we calculate the difference between the time of maximal peak of rotation and the time of maximal peak of translation, namely, t(R*)-t(T*). This procedure shows that translation precedes rotation before immobility (I), and that rotation precedes translation after immobility (Fig. 4C). While there is high variability in the time intervals across animals, all follow the same sequential order of global peaks.t(R*)-t(T*)

**Figure 4.**
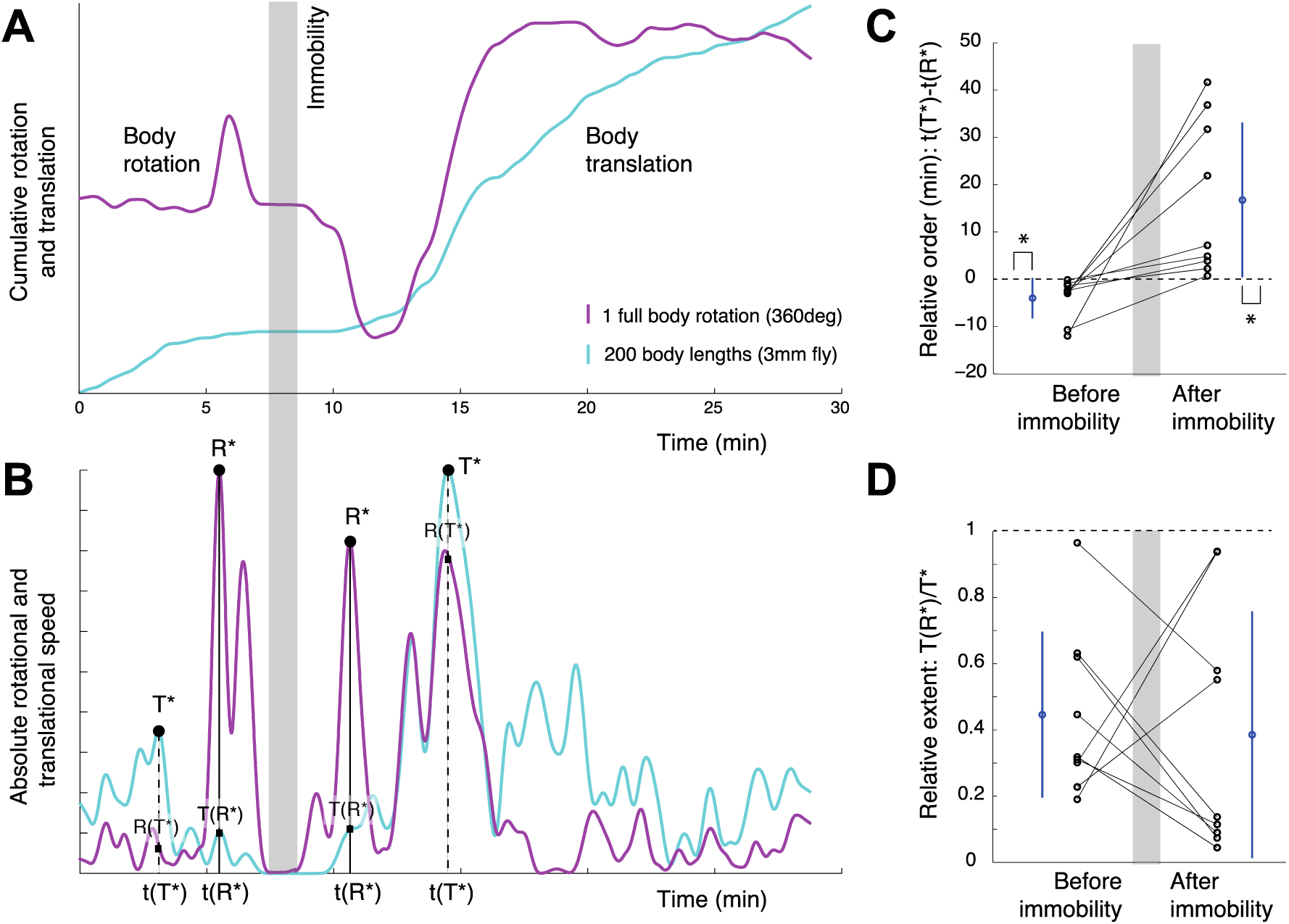
Synchronic and diachronic dynamics of translation and rotation. (**A**) Cumulative body rotation (magenta) and translation (cyan) reveal the global sequence of changes in the rotational and translational degrees of freedom. The shaded area marks the period of immobility, which is used as a reference for measuring the events that precede and follow it. (**B**) Global changes in speed of progression and body orientation are obtained from the absolute time derivative of the curves in (A). As shown, a global peak in translation followed by a global peak in rotation precede immobility, and a global peak in rotation followed by a global peak in translation follow immobility (TRIRT sequence). (**C**) Difference between the time of maximal peak of translation and the time of maximal peak of rotation, t(T*)-t(R*), is negative for transitions into immobility (p=0.004, Sign test) and positive for transitions out of immobility (p=0.004, Sign test). Absolute time differences are greater for transitions out of immobility than for transitions into immobility (p=0.0352, Sign test). Each line connecting dots represents the same animal. Mean and standard deviation in blue. (**D**) Quantification of the strength of reciprocity in the value of global peaks of rotation and translation during transitions in and out of immobility show hardly any difference. Dots represent the score for individual flies and lines connect the results for the same individual. The mean and standard deviation are colored in blue.

Next, in order to quantify the relative strength of the reciprocal relationship between global peaks, we calculate the value of translation at its peak, T(T*), and compare it with translation when rotation is at its peak T(R*). Thus we measure the amount of reduction in translation by the time that rotation reaches its peak. The smaller the ratio T(R*)/T(T*), the stronger the phenomenon of reciprocity between translation and rotation (Fig. 4D). Note that this relationship is by no means reciprocal at all times: in the presented graph there are cases in which both rotation and translation increase, and also in which both decrease together. In other words, rotation and translation are globally, *not* locally, reciprocal. Characterizing the relative strength of rotation and translation peaks by means of the above ratio is invariant to time rescaling and to absolute values of rotation and translation. This is necessary for capturing the invariance in the sequence and strength across individual animals, and particularly useful given the large variability in the timescales of unfolding of the phenomenon (some flies take minutes, others take hours) and in the rotation values (some flies perform ten full body rotations, others perform hundreds; and they do so at different rates).

On the whole, flies follow the same sequence of transition into immobility, involving, for each dimension separately, an enhancement, a reduction, and then elimination of that degree of freedom, thus progressively narrowing down the fly’s locomotor repertoire; and the same but opposite sequence of transition out of immobility, involving, for each dimension separately, an enhancement, and then subsiding to normal of that degree of freedom, thus progressively building up the fly’s locomotor repertoire. Such invariance can be summarized by the acronym TRIRT (generative rule consisting on the sequence: Translation, Rotation, Immobility, Rotation, and Translation).

### Rotational switching rate changes in and out of immobility

One predominant effect of cocaine administration is the high rate of repetition of full body rotations and their rotational speed. During rotation the animal may switch between clockwise (CW) and counterclockwise (CCW) rotations. It is now possible to examine the dynamics of switching in reference to immobility, in the context of the animal’s freedom of movement. To do so, we focus on how frequently the fly changes the direction of rotation and how biased successive rotations are. Globally, there are long-term predominant biases to rotate in a particular direction (Fig. 5A). In particular, transitions out of immobility start by very long rotations in the same overall direction. However, flies do not rotate monotonically in one direction but, rather, alternate between large amplitude rotations in one preferred direction and low amplitude rotations in the other direction (Fig. 5B-C). Locally, the fly alternates between CW and CCW rotations. The switching rate decreases when transitioning into immobility and increases when transitioning out of it (Fig. 5D-F). As found for the synchronic relationship between translation and rotation (TRIRT), this diachronic pattern for rotational switching also exhibits a mirror symmetry between the process leading into immobility and out of it (Fig. 5E-F), and can be summarized by the acronym SsIsS (sequence of high Switching, reduced switching, Immobility, and the reverse).

**Figure 5.**
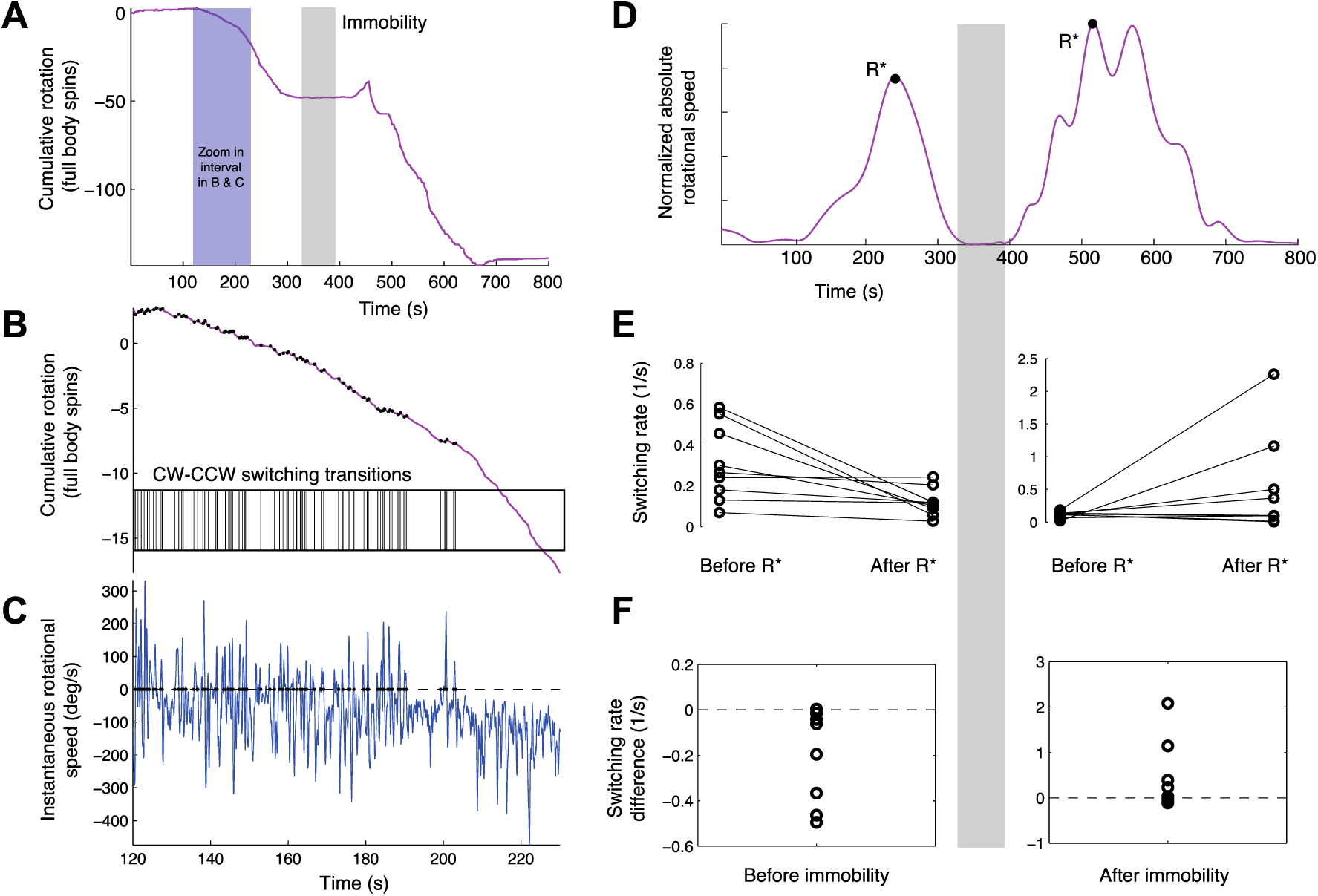
Switching between clockwise and counterclockwise rotation decreases into immobility and increases out of immobility. (**A**) Cumulative body orientation as a function of time for a single fly across the session. The gray shaded area marks the period of immobility. The blue shaded area marks a zoomed-in time interval presented in (B) and (C). As shown, the general tendency of this fly is to rotate clockwise both before and after immobility. Note that during transition into immobility the fly performs in the order of 50 full body rotations in 5 minutes, and during transition out of immobility, as many as 100 full body rotations in 5 minutes. (**B**) Zoomed in time segment in (A) of cumulative body rotation. As illustrated, while the fly generally rotates clockwise, it keeps switching between clockwise and counterclockwise rotations. Each transition is marked by a small black dot on the curve and a corresponding vertical bar on the transitions raster plot below. (**C**) Time derivative of the fly’s momentary body orientation. Zero crossings in the rotational speed mark switching, corresponding to the vertical bars in (B). As shown, there is an overall drift in one direction, concurrent with a decrease in the number of transitions as a function of time. (**D**) Global changes in body orientation obtained from the absolute time derivative of the curve in (A). To examine the change in switching across the session, the period leading to immobility is partitioned into the segments preceding and following maximal rotation (same for the period leading out of immobility). (**E**) Switching rate, calculated as the number of transitions per second, decreases in the interval preceding immobility (left panel, p<0.039, Sign test) and tends to increase in the interval following immobility (right panel, not significant). Dots represent the score for individual flies and lines connect the results for the same individual. (**F**) Overall the difference of the switching differences after immobility and before it is always positive (p<0.00195, sign test) and has a median of .37 switches per second. Dots represent the switching rate change for every fly.

## Discussion

### Generative rules shaping fly cocaine-induced behavior during transitions out and into immobility

Using immobility as a reference for the measurement of behavior and cocaine as the parameter inducing a behavioral gradient, we found that the flies exhibit a progressive buildup of their locomotor repertoire when starting from immobility, and a progressive narrowing down of the repertoire towards into immobility. During buildup, for each key variable separately, the fly exhibits first enhancement and then reduction to normal values of movement along that variable: first, of body rotation in the horizontal plane, then of path curvature, and then of speed of translation. The extents of movement across the key variables show reciprocal relations: when rotation is at its peak translation is low, and when translation is at its peak rotation is low, while path curvature is partly coupled to rotation. Transition into immobility from rich normal locomotor behavior unfolds through narrowing down of the repertoire in the opposite sequential order, also showing reciprocal relations between the extents of the same variables. Quantification of the generative rules of this behavior, based on the temporal sequence of global peaks of extent (Fig. 4) provides a summary of the *bauplan* of this arthropod behavior, allowing a comparison with the rules reported in previous studies for movement into and out of immobility in vertebrates, in which the behavior has been termed The Mobility Gradient^9^.

### Vertebrates and fruit flies share the same generative rules

*Buildup:* In infant mice transition out of novelty-induced immobility consists of side-to-side head movements that increase in amplitude, gradually recruiting the forequarters and then the hindquarters, to extensive rotation in place around the hindquarters. Only after exhausting the horizontal plane by rotating around the hindquarters, does forward stretching and subsequently forward translation appear, first along curved and then along straight paths (movie S5). The same sequence is exhibited in amphibians^15^, fish^16^, insectivores (movie S6), and carnivores^17^. Head-raising, forequarter-raising and, finally, rearing on the hind legs, are exhibited next^18,19^. The same sequence is exhibited both during moment-to-moment behavior and in ontogeny, during recovery from lateral hypothalamic damage^20^. Later on in development, during, for example, play and ritualized fighting interactions, the inferior animal exhibits the less mobile portion of the sequence, culminating in rearing and rotating around the hindquarters, whereas the superior may rear and rotate both around the hind-legs and around the forelegs, exhibiting an expanded freedom of movement in both the horizontal and vertical dimensions^17,21^; for a review see^9,22^ (movie S7).

*Narrowing down*: The opposite sequence, proceeding from rich normal behavior to enhanced, then reduced, then immobility, first of rearing, then of translation along straight, and then along curved paths, then of rotation, culminating in relative immobility, is exhibited in rats following the administration of several dopamine agonists^4,5,23-27^ (movie S8). This mobility gradient^9,28^, which is composed of buildup (warmup) and narrowing down (shutdown) sequences^18,19^ shares with the mobility gradient demonstrated in fruit flies the same origin (immobility), parameter (dopaminergic stimulation; but in vertebrates also novelty and proximity to a rival), and generative rules.

As there are no “fossil skeletons" of behavior we suspend judgement regarding common descent. From the vantage point of comparative anatomy, generative rules define the core, or skeleton that has resisted adaptation and around which there is a variable adaptive component. As shown by ethology, this core/adaptive distinction also applies to behavior^29,30^. The shared generative rules of the mobility gradient that we expose constitute a dynamical version of the “principle of connections” of St. Hillaire (“equivalence under transformation”^2,7,8,31^) whereby homology must be defined, not by its form, but by the relative positions, spatial, and, one might add, temporal relations between the elements of a structure.

Applying “serial homology” when same structure serves different functions in the same species

Using our *generative rules* as a search image, not only can we predict essential sameness in the brain/behavior interface of both flies and rodents, but we can also anticipate that the same rules might underlie apparently different functional behaviors in the same species. This is similar to identifying serial homology in anatomy, where, e.g., hand and foot are considered homologous because they share the same set of developmental constraints, caused by locally acting self-regulatory mechanisms of differentiation^32^. Reviewing the fruit fly larval behavior literature with a search image for low and high mobility, attention is immediately drawn to the abnormally high extent of turning behavior exhibited by larvae with mutations in the gene *scribbler* (sbb) in the absence of food^33^. These appear to be respective manifestations of the high and low ends of the mobility gradient. The four key features characterizing low mobility in the cocaine-treated fly (low speed of translation, highly curved path, high body rotation, and immobility) exhibit a full correspondence to the features of the “abnormal crawling pattern” exhibited by *scribbler* larvae: low speed, curved paths, high turning rate, and long pauses^34^. The parameter precipitating this behavior could be, as Sokolowski and co-workers suggest, the absence of food, or else, given our search image, the stress brought about by the absence of food, or even its presence in hungry flies^35,36^. Equivalent differences in mobility, expressed by pivoting and/or rearing on hind legs and forelegs, reported to be exhibited by vertebrate partners engaged in interactions ^17,21,28^, might also be looked for in fruit fly courtship and agonistic interactions.

Searching for equivalent neurochemical substrates mediating the buildup and narrowing down

Recently, it has been claimed that the vertebrate basal ganglia and the arthropod central complex are deeply homologous. In both, comparable systems of dopaminergic neurons, their projections, and dopaminergic receptor activities are involved in the modulation and maintenance of behavior^37^. Dopamine systems are also key players in generating and regulating the mobility gradient ^12,13,23-27^, and dopaminergic stimulation of specific substructures of the basal ganglia and central complex induce specific components of the mobility gradient respectively in rodents ^38-42^ and in flies^43^. The neurochemical processes mediating the buildup of the vertebrate locomotor repertoire have been recently attributed by Cools and co-workers to dopaminergic feedforward loops operating in the basal ganglia^12,13^. The equivalence between the mobility gradient core phenomena and the feedforward loops exposed in the basal ganglia can be used as a search image or working hypothesis for studying the relations between the arthropod mobility gradient and the central complex. It might be useful to examine whether feedforward loops also mediate the buildup of locomotor behavior functions in the central complex.

Given the foundational role of behavior in neuroscience^53^, it is perhaps time to start relating the various levels of the biological hierarchy to the behavioral *bauplan* they support. Here, practicing the methodology of comparative anatomy to the study of behavior, we have established a shared behavioral architecture across distant phyla based on the principle of connections. Having established the common behavioral architecture of the mobility gradient in the two phyla, demonstrating that they are respectively mediated by deeply homologous neural structures would endow the Mobility Gradient with the status of a Darwinian homology.

## Materials and Methods

### Fly stocks

Drosophila cultures were maintained at 24°C on a standard cornmeal-molasses medium in 12 hour light-dark cycle at 60% humidity. The experiments were performed on three-day-old flies of the wild-type laboratory strain Canton-S. Nine male flies were tested in a low-throughput high-content-data approach.

### Behavioral arena

The experimental setup for observing and tracking the flies was a 15 cm diameter circular arena with 0.7cm height wall and glass ceiling. The arena was illuminated from above with a 40W bulb. A camera placed above the arena recorded the fly’s behavior. During the experiment there was a continuous airflow through the arena, through two small wall openings allowing also the introduction of the volatilized drug into the arena during the experiment^14^.

### Animal preparation

Neither food nor water was supplied to the fly during the entire experiment. All experiments were performed during the 12 hours light period, and on one fly at a time. Each fly was transferred to the arena and allowed to habituate there for one hour. Upon exposure to cocaine, the fly behavior was uninterruptedly recorded including full sedation (immobility) and the process of recovery (regaining normal locomotor behavior).

### Behavioral tracking

The fly locomotor behavior was recorded at 25 frames per second using a CCD camera. Following video acquisition, the centroid position of the fly and its body axis direction were tracked with FTrack^44^, a custom-made software written in Matlab (Mathworks). Raw trajectory data were corrected for tilt and rotation of the camera. Data segments during which it was not possible to assess the fly’s orientation (fly located on the wall or jumping) were excluded from analysis.

### Behavioral analysis

Quantitative analysis of the animal’s behavior was based on the dynamics of three main degrees of freedom: centroid speed, path curvature and body rotation. Measuring the direction of progression and the body orientation allows to distinguish where the animal is going versus where it is facing^14,46,47^ (note that the fly can walk north and then turn left, while still having the freedom to face north). In the Eshkol Wachman Movement Notation (EW) these variables amount to speed and direction of shift of weight, and direction of front^45^. Changes in the direction of progression are calculated per unit of progression and as a function of time in order to have a geometric curvature^48,49^ expressed in kinematic terms^50,51^. Switching between clockwise and counterclockwise body rotation was assessed via the zero-crossings of the instantaneous time derivative of the body angle, removing artifacts during arrests by pruning out rotations smaller than 12 degrees. Since we studied the same phenomenon across a wide range of timescales (from subsecond small oscillations during rotation in place, to hour-long sedation recovery), we produced reliable estimates of local and global variables in time by using the variable-window smoothing LOWESS method^52^. Immobility, defined as the longest time interval of complete arrest (no translation nor rotation) across the whole session, allows to transform chronological time into activity, revealing dynamical invariants despite animal-to-animal behavioral variability.

## Acknowledgements

This study was funded by the Portuguese Foundation for Science and Technology (FCT grant No SFRH/BPD/97544/2013 to AGM) and by the Israel Science Foundation (grant No 760/08 to IG and YB). We thank the participants of the “Homology in NeuroEthology” course held at the Champalimaud Neuroscience Programme for fruitful discussions. We acknowledge the gifts of fly stocks from Dani Segal. We thank Gonçalo Lopes, Eduardo Dias-Ferreira, Andre Brown, Lauren McElvain, Troy Shirangi, Eyal Gruntman and Ehud Fonio for valuable feedback on the manuscript.

## Author contributions

I.G., Y.B., and A.G.M. conceived the research; E.O. performed the experiments; D.V. developed the tracking software; A.G. preprocessed the data; Y.B. and A.G.M designed the quantitative analysis; A.G.M. analyzed the data and made the figures; A.G.M, Y.B, and I.G. wrote the paper.

## Supplementary Movies

**movie S1**: https://www.youtube.com/watch?v=bgRyVv8Ae04

**movie S2**: https://www.youtube.com/watch?v=LBAugS1Rk7A

**movie S3**: https://youtu.be/CzT7sn0RdkM

**movie S4**: https://youtu.be/9vkTYuFQLSI

**movie S5**: https://www.youtube.com/watch?v=wVaeqWPZnfc

**movie S6**: https://www.youtube.com/watch?v=3JQZgTwQrwM

**movie S7**: https://www.youtube.com/watch?v=YzD21jpa09g

**movie S8**: https://www.youtube.com/watch?v=ittpzspGZ_g

## Supplementary Figures

**Figure S1.**
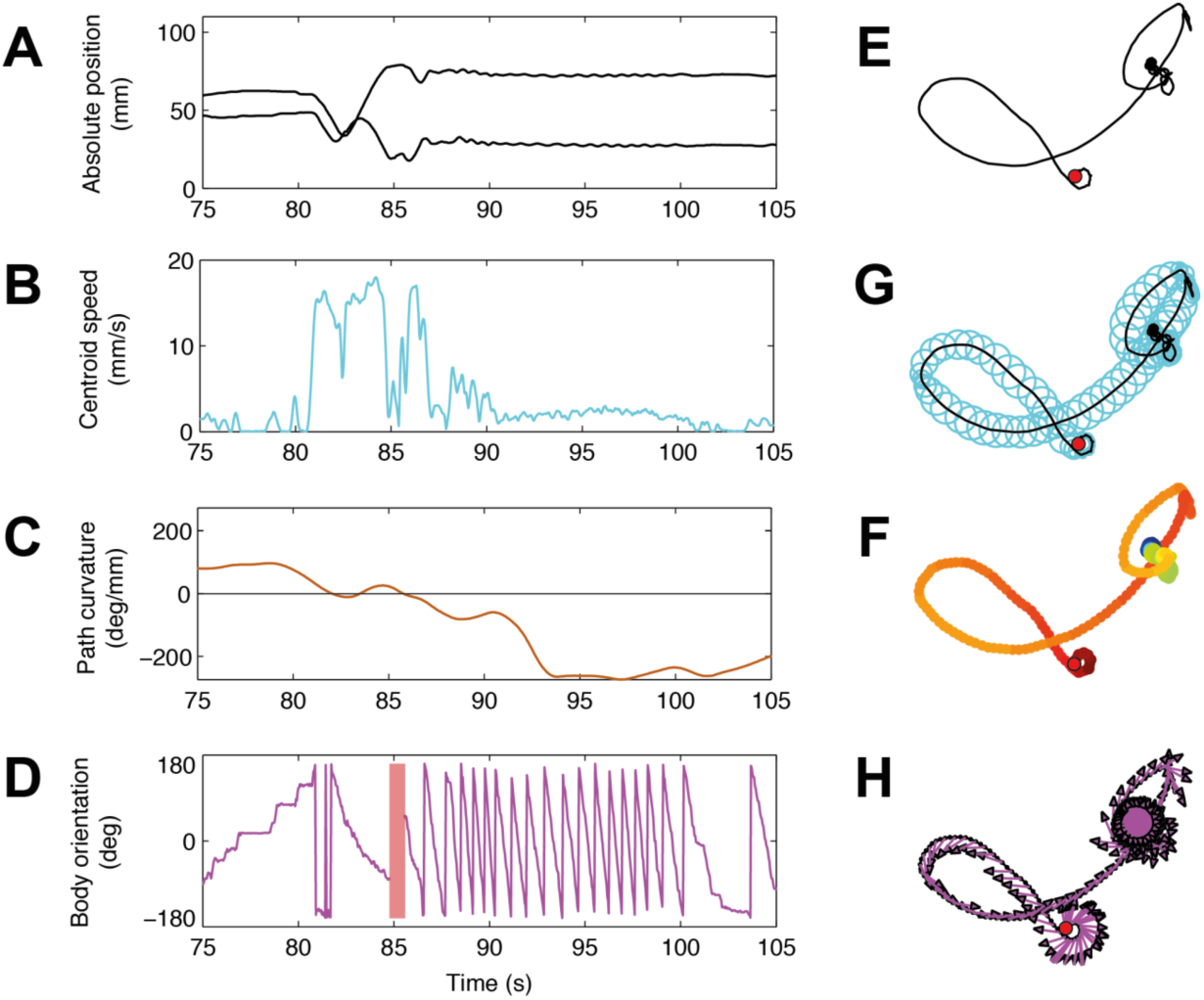
Three main kinematic degrees of freedom exercised by a fly at the trajectory level. Time evolution of XY position (**A**), centroid speed (**B**), path curvature (**C**), and body orientation (**D**). Same degrees of freedom presented in space: black line represents fly path (**E**), disk diameter is proportional to speed (**G**), color coding depicts curvature (**F**), and arrows depict body orientation (**H**). Small red dot marks the beginning of the trajectory. During the 30 second time segment illustrated, a burst in speed as the animal traces a relatively straight path is followed by low speed at very high curvature, while the animal vigorously rotates in place, at approximately 360 degrees per second. Taking into account the orientation of the longitudinal axis of the fly’s body as distinct from the direction of progression, we can access a third degree of freedom, thus distinguishing and quantifying in what direction the animal is moving, how fast it is progressing, and in what direction it is facing.

**Figure S2.**
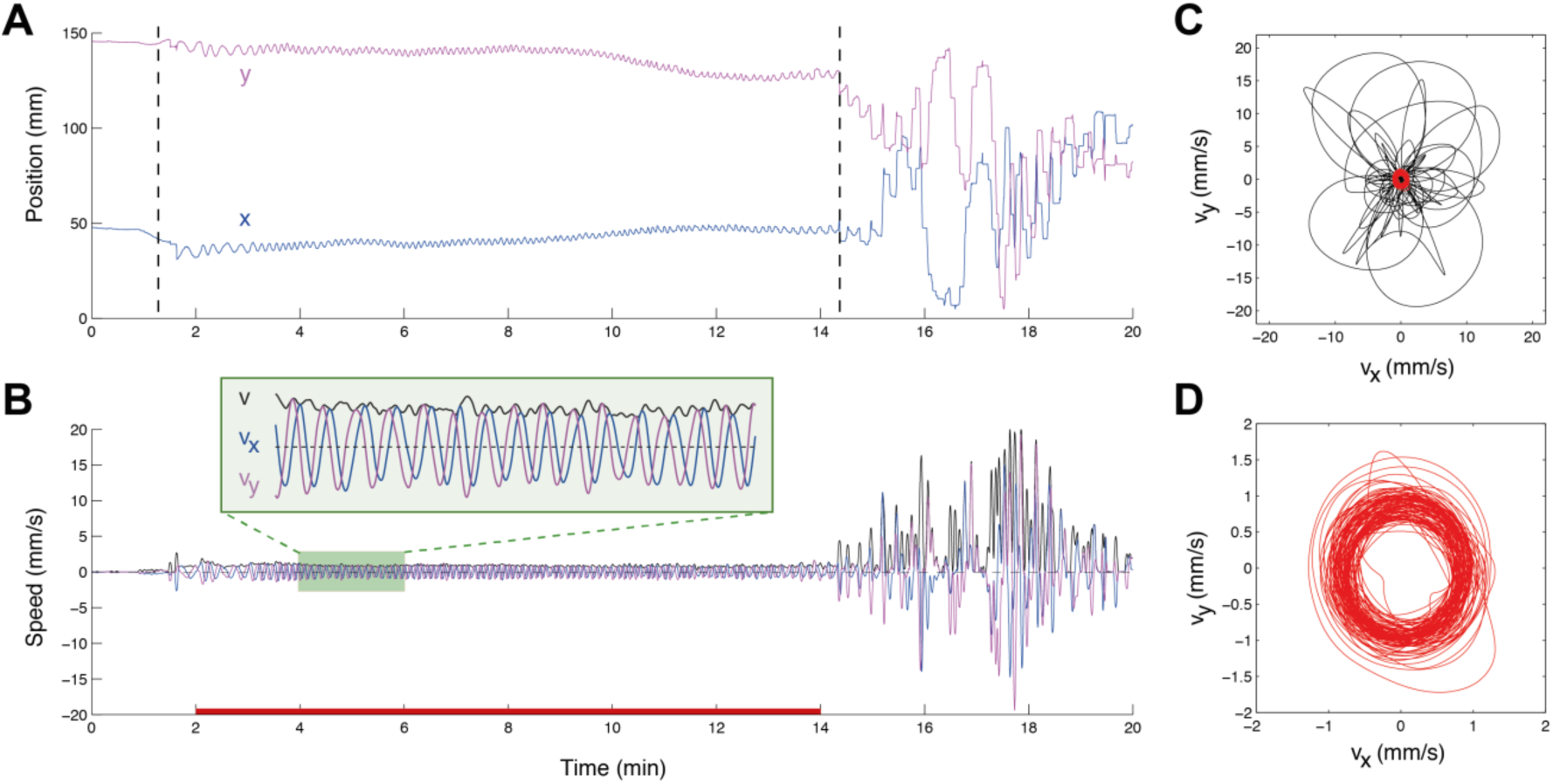
Detailed dynamics of the transition out of immobility based on velocity components, which determine the dynamics and coordination of speed and curvature. (**A**) Transition out of immobility is illustrated by plotting the x and y positions as a function of time. Starting from absolute immobility, the fly performs tiny but fast oscillations in the x and y positions, reflecting fast rotations in place, which progressively slow down, and finally change to large displacements, corresponding to normal progression. (**B**) The early stage of transition out of immobility is characterized by very low speeds, whose perpendicular components (vx and vy) alternate in oscillatory dynamics (vx is zero when vy is max, and vice-versa; see inset in green), corresponding to very high curvature. The velocity components along the x and y directions show the coordinated circling as the fly transitions out of immobility, whereas speed (v) does not capture the subtleties of rotation in place. (**C**) Phase-plot of speeds along x and y directions, containing both low speeds (in red), but also high-speed progression segments in all directions at a later stage of transition out of immobility. (**D**) Zoom in of the plot in (C) showing only the velocity components during the time interval from minute 2 to minute 14. On the whole, this closer look at path dynamics reveals that high curvature emerges from small, fast and alternating oscillations in orthogonal components of the velocity vector at typical speeds of 1mm/s, which reveal the minute circles traced by the animal as it rotates in place. During walking at higher speeds, flies show a range of different velocity components.

